# Single cell visualisation of *mir-9a* and *Senseless* co-expression during *Drosophila melanogaster* embryonic and larval peripheral nervous system development

**DOI:** 10.1101/2020.07.27.223479

**Authors:** Lorenzo Gallicchio, Sam Griffiths-Jones, Matthew Ronshaugen

## Abstract

The *Drosophila melanogaster* peripheral nervous system (PNS) comprises the sensory organs that allow the fly to detect environmental factors such as temperature and pressure. PNS development is a highly specified process where each sensilla originates from a single sensory organ precursor (SOP) cell. One of the major genetic orchestrators of PNS development is *Senseless*, which encodes a zinc finger transcription factor (Sens). Sens is both necessary and sufficient for SOP differentiation. *Senseless* expression and SOP number are regulated by the microRNA *miR-9a*. However, the reciprocal dynamics of *Senseless* and *miR-9a* are still obscure. By coupling smFISH with immunofluorescence, we are able to visualize transcription of the *mir-9a* locus and expression of Sens simultaneously. During embryogenesis, we show that the expression of *mir-9a* in SOP cells is rapidly lost as *Senseless* expression increases. However, this mutually exclusive expression pattern is not observed in the third instar imaginal wing disc, where some *Senseless*-expressing cells show active sites of *mir-9a* transcription. These data challenge and extend previous models of *Senseless* regulation, and show complex co-expression dynamics between *mir-9a* and *Senseless*. The differences in this dynamic relationship between embryonic and larval PNS development suggest a possible switch in *miR-9a* function. Our work brings single-cell resolution to the understanding of dynamic regulation of PNS development by *Senseless* and *miR-9a*.

## Introduction

One of the most impressive demonstrations of developmental robustness is the specification of the *Drosophila melanogaster* peripheral nervous system (PNS), which comprises all the organs that allow the fly to detect movement, pressure, temperature, and more. *Drosophila* sensilla number and position exhibit little or no variation from individual to individual, even in diverse environmental conditions (Hartenstein 1988). The key early developmental step involves the selection and specification of sensory organ precursor (SOP) cells from a field of equipotent cells. During early embryogenesis (~5h from fertilization), groups of epidermal cells start to express Achete-Scute complex genes. These proneural genes impart the potential to become neurons (Jarman *et al*. 1993; Jan and Jan 1994; Goulding *et al*. 2000; Huang *et al*. 2000; Reeves and Posakony 2005). This potential is then constrained via Notch lateral inhibition to a single cell in the cluster, the SOP cell (Ghysen and Dambly-Chaudiere 1989; Artavanis-Tsakonas and Simpson 1991). The many different classes of sensory organs all originate from SOPs and develop via common shared rounds of cellular division (Lai and Orgogozo 2004). The eventual differences between sensory organs arise in part through subsequent changes in cell death and proliferation (Orgogozo *et al*. 2001; Orgogozo and Schweisguth 2004).

One of the major effectors of PNS development is a gene named *Senseless* (*Sens*) (Nolo *et al*. 2000). *Sens* encodes a transcription factor (Sens) whose expression is initially activated and subsequently maintained by the proneural genes *achete* and *scute* (Jafar-Nejad *et al*. 2006). *Sens* in turn maintains the expression of proneural genes to direct proper neuronal cell differentiation (Nolo *et al*. 2000; Acar *et al*. 2006). *Sens* expression is first detectable during stage 10 of *Drosophila* embryogenesis, as isolated cells start to specify according to their SOP fate potential. As embryogenesis proceeds, these isolated *Sens*-expressing SOPs ultimately give rise to the entire sensory organ. *Sens* expression becomes repressed around stage 13, when SOPs are fully specified (Nolo *et al*. 2000). Since *Sens* maintains proneural gene activation, loss-of-function *Sens* mutant embryos exhibit a decreased number of SOPs, corresponding to a loss of sensory organs in the adult fly (Nolo *et al*. 2000). Gain of function mutations and ectopic expression of Sens cause an increased number of SOPs and consequently sensory organs (Jafar-Nejad *et al*. 2003; Li *et al*. 2006). Therefore, it is suggested that *Sens* is necessary and sufficient for SOP differentiation (Nolo *et al*. 2000). The robustness and reproducibility of sensory organ development between individuals implicates *Sens* as a keystone gene whose fine-scale regulation involves multiple feedback inputs.

Neurogenesis is extensively regulated by microRNAs (miRNAs) (Nolo *et al*. 2000; Hilgers *et al*. 2010; Caygill and Brand 2017). These small regulators of translation and mRNA stability contribute to the robustness of many biological processes. It has been shown that *miR-263a/b* stabilize sensory organ patterning in the retina by inhibiting SO cell apoptosis (Hilgers *et al*. 2010), and that *miR-7* stabilizes neuronal differentiation in the *Drosophila* larval brain by targeting the Notch pathway (Caygill and Brand 2017). In addition, Li and co-workers (2006) showed that *miR-9a* regulates Sens function through multiple target recognition sites in the Sens 3’ UTR. When Sens’ *miR-9a* binding sites are mutated, Sens levels are not only higher but more sensitive to temperature perturbations (Cassidy *et al*. 2013), resulting in an altered distribution of sensory organs in the wing margin (Cassidy *et al*. 2013; Giri *et al*. 2020). Loss-of-function and over-expression of *miR-9a* produce opposite phenotypes with respect to *Senseless* in both embryos and larvae. Thus the phenotypic consequences of *miR-9a* disruption mirror those of *Sens*, suggesting that *miR-9a* is necessary to ensure appropriate Sens expression in the right cells and at the right level to convey robustness to SOP specification (Li *et al*. 2006).

The *miR-9a* microRNA is a member of one of the ~30-40 families that are predicted to pre-date the divergence of protostomes and deuterostomes, and therefore to be conserved in essentially all bilaterian animals (Wheeler *et al*. 2009; Ninova *et al*. 2014). In every animal where *miR-9* family members have been studied functionally, they are found to regulate processes related to neurogenesis and neuronal progenitor proliferation (Sempere *et al*. 2004; Wheeler *et al*. 2009; Delaloy *et al*. 2010). For instance, over-expression of *miR-9* in zebrafish embryo (Leucht *et al*. 2008), mouse embryonic cortex (Zhao *et al*. 2009) and chicken spinal cord (Yoo *et al*. 2011) all lead to a reduction of the number of proliferating neural progenitors and impairment of PNS development. These studies demonstrate clear similarities between *miR-9* expression and function in *Drosophila* and vertebrates. Disrupted *miR-9* function has also been linked with some human pathologies, including cancer progression (Nowek *et al*. 2018) and neurodegenerative amyloid diseases (Packer *et al*. 2008). For instance tumorigenic cells from medulloblastoma appear to have decreased expression of *miR-9*, while a subclass of glioblastoma tumour cells express *miR-9* at a higher level (Ferretti *et al*. 2009; Kim *et al*. 2011). In addition, *miR-9* has been also found to have a role as a proto-oncogene and/or as a tumour-suppressor gene during progression of cancers not directly related with the nervous system (Coolen *et al*. 2013).

The current model of *miR-9a* function in *Drosophila* SOP specification suggests mutually exclusive reciprocal expression of *miR-9a* and *Sens* in SOPs (Li *et al*. 2006). This in turn suggests a role for *miR-9a* in ensuring that only one of the cells in the progenitor field takes on SOP identity. In this work, we use single-cell quantitative FISH and nascent transcript FISH to investigate the *miR-9a/Sens/SOP* regulatory model in hitherto unseen detail. This use of single-molecule *in situ* hybridization (smFISH) coupled with immunofluorescence (IF) allows us to simultaneously visualize active sites of *miR-9a* transcription and Sens protein in both embryos and larval wing disc at the single cell level. We use these data to analyse the dynamics of *miR-9a* transcription and Sens protein abundance. We find that *miR-9a* and Sens are initially co-expressed, but ultimately exhibit a dynamic reciprocal expression pattern. We observe that *miR-9a* transcription becomes rapidly repressed in high Sens expressing SOPs during embryogenesis, presumably as Sens protein accumulates in the cell nucleus. A subtly different co-expression dynamic was observed during wing disc development, where many SOPs also express *miR-9a*. These SOPs exhibit an inverse relationship between Sens abundance and *miR-9a* transcription. These new data refine and expand the previous model to provide key new insights into *miR-9a/Sens* regulation in PNS development (Li *et al*. 2006). In particular, we include for the first time a temporal element to the understanding of the dynamics of *miR-9a* regulation of SOP differentiation.

## Methods

### Fly stocks, embryo collection and fixing and larval dissection

Flies were grown at 25°C or 18°C. Embryos were collected after ~20h and fixed in 1V heptane + 1V 4% formaldehyde for 30 minutes shaking at 220 rpm. The embryos were then washed and shaken vigorously for one minute in 100% methanol. Fixed embryos were stored in methanol at −20°C. Larvae were dissected in 1XPBS, carcasses were fixed in 1V 1XPBS + 1V 10% formaldehyde for ~1h, washed with methanol and stored in methanol at −20°C.

Genotypes used for this study are: W [1118], Bloomington stock 3605 and 2XTY1-SGFP-V5-preTEV-BLRP-3XFLAG-Sens, VDRC stock ID 318017

### Probe design, smFISH and immunofluorescence

We adapted the inexpensive version (Tsanov *et al*. 2016) of the conventional smFISH protocol in *Drosophila* (Trcek *et al*. 2017). Primary probes were designed against the mature *Senseless* and sfGFP mRNA and a genomic region flanking the *mir-9a* gene locus, all from FlyBase, using the Biosearch Technologies Stellaris probe Designer (version 4.2). To the 5’ end of each probe was added the Flap sequence CCTCCTAAGTTTCGAGCTGGACTCAGTG. Multiple secondary probes that are complementary to the Flap sequence were tagged with fluorophores (CAL Fluor Orange 560, CAL Fluor Red 610, Quasar 670) to allow multiplexing.

Antibodies used were Anti Green Fluorescent Protein rabbit igG fraction (Invitrogen #A11122) at 1:500, anti-Sens (Nolo *et al*. 2000) at 1:1000, Goat anti-Guinea Pig IgG (H+L) Highly Cross-Adsorbed Secondary Antibody Alexa Fluor 555 (Invitrogen #A21435) at 1:500, and Goat anti-Rabbit IgG (H+L) Cross-Adsorbed Secondary Antibody Alexa Fluor 488 (Invitrogen #A11008) at 1:500.

### Imaging and quantification

Imaging was performed using a Leica SP8 Inverted Tandem Head confocal microscope with LAS X v.3.5.1.18803 software (University of Manchester Bioimaging facility), using 20X, 40X and 100X magnifications. Deconvolution was performed using Huygens Pro v16.05 software. Protein fluorescence levels were measured using FIJI for Macintosh. From each picture, five measurements of background mean intensity were taken. Each single cell measurement was then adjusted using the formula: integrated density of nucleus – (area of nucleus x background mean).

## Results

### *miR-9a* is expressed in the dorsal ectoderm during embryogenesis and ubiquitously in the wing disc

In order to understand the interaction between *miR-9a* and its target *Senseless*, we first systematically describe the *mir-9a* expression pattern in the embryo and imaginal disc at the single cell level. Imaging mature miRNAs is difficult. Previously applied techniques require amplification and often have significant background noise problems (e.g. probes labelled with digoxigenin) (Biryukova *et al*. 2009). Many researchers have tried to overcome this (Lu and Tsourkas 2009), but these approaches still have very limited use for single cell analysis and quantification.

We have used a nascent transcript approach using single molecule FISH to detect expression in cells that are actively producing the miRNA primary transcript (pri-miRNA). To do so we have designed sets of ~48 probes (table 3) that hybridize to a ~1kb region in the primary miRNA transcript flanking the *mir-9a* locus. This allows the detection of active *mir-9a* transcription in the cell nuclei as previously described by Aboobaker and co-workers (2005). We find that expression of *mir-9a* initiates in early stage 5, at which point it is expressed throughout the dorsal ectoderm of the embryo (Fig. 1 A-A’). The pattern displays a precise boundary between actively transcribing cells and non-expressing cells, which correlates with the mesoderm-ectoderm boundary similar to that described by Fu et al. (2014). There are also small domains at the posterior and anterior embryonic ends lacking *mir-9a* expression. Later, during *Drosophila* embryonic stages 6 and 7, *mir-9a* expression is maintained in this pattern throughout the ectoderm (Fig. 1 B-B’ and Fig.1 C-C’), clearly marking the boundary between ectodermal cells and invaginating mesodermal cells. At stage 8, the mesoderm is almost completely invaginated and the *mir-9a* expression domain covers the embryo surface (Fig. D-D’), with the exclusion of the same regions at the anterior and posterior ends. At stages 11 and 14, *mir-9a* continues to be expressed throughout the ectoderm, except from a dorsal anterior region, and it is largely absent from the amnioserosa (Fig. 1 E-F).

**Figure 1.**
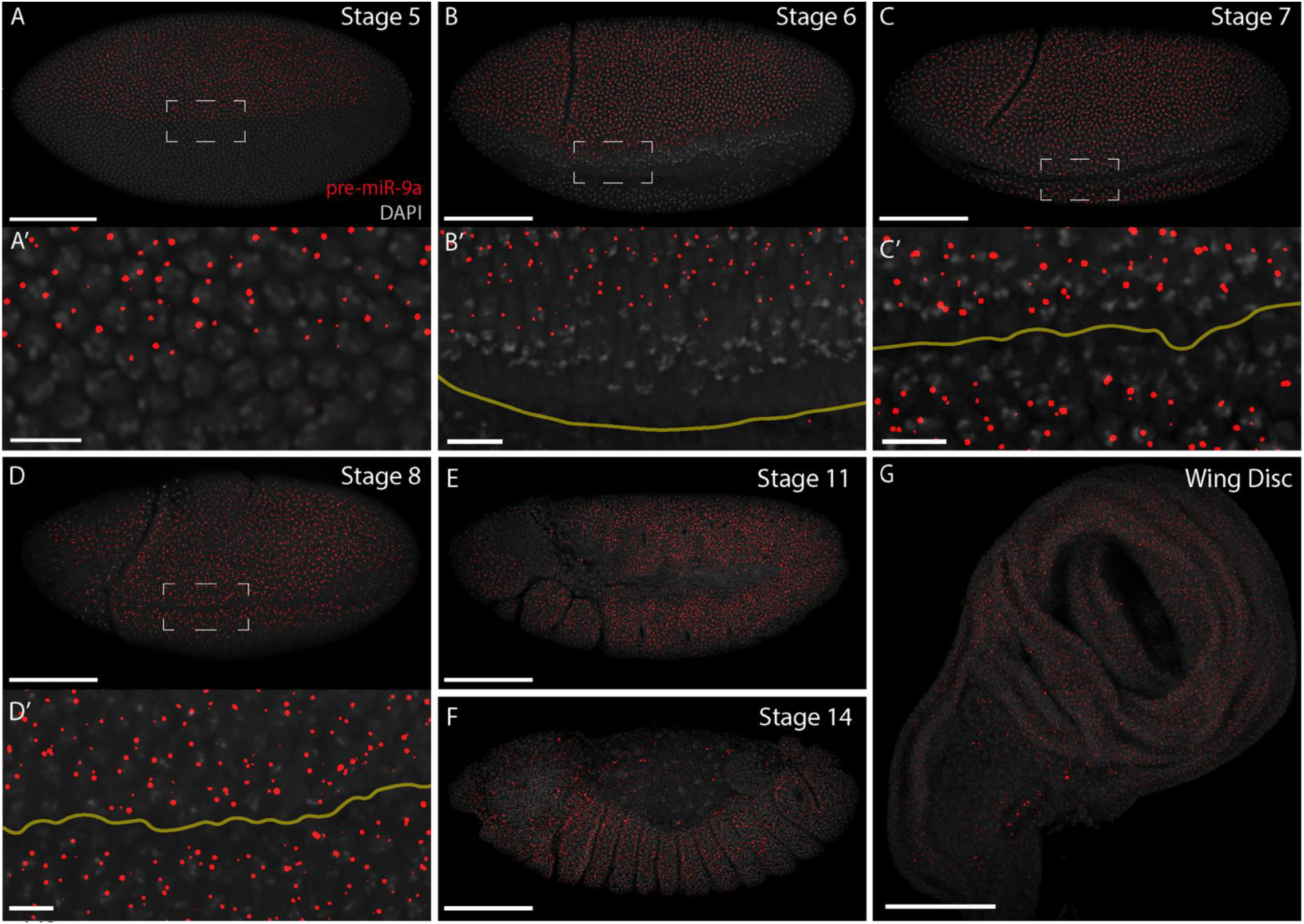
*mir-9a* expression pattern during embryogenesis and in the wing disc. Cells that are actively transcribing *mir-9a* present one or two puncta, indicating *mir-9a* active TSSs. (A-A’) Stage 5 WT *Drosophila* embryo: *mir-9a* is expressed in the dorsal ectoderm, before the ventral furrow is evident. (B-B’) Stage 6 WT: The *mir-9a* expression domain extends towards the forming ventral furrow (highlighted with a yellow line). (C-C’) Stage 7 WT. (D-D’) Stage 8 WT. (E-F) At later stages, *mir-9a* is expressed throughout the ectoderm. (G) WT *Drosophila* wing disc: *mir-9a* is expressed widely throughout the disc. Scalebars: (A) 100 μm, (A’) 10 μm, (G) 100 μm.

It has previously been reported that *mir-9a* is expressed widely in the 3^rd^ instar wing disc but not in cells expressing *Senseless* (Li *et al*. 2006; Biryukova *et al*. 2009). Similarly, we observed that *mir-9a* is actively transcribed everywhere in the wing disc, with the exception of a few randomly distributed cells (Fig. 1 G).

### Immunodetection of *Sens*-sfGFP fusion protein allows the study of *Sens* expression in *Drosophila* embryos at single-cell resolution

The role of *miR-9a* in regulating the transcription factor *Senseless* (*Sens*) is well characterised genetically during SOP specification (Li *et al*. 2006; Cassidy *et al*. 2013). To investigate the dynamics of *miR-9a* regulation of *Sens* we developed a strategy to simultaneously observe *Sens* transcription and protein accumulation at the single-cell level via confocal fluorescent microscopy. In principle, the efficient detection of proteins in fixed samples using IF relies on the availability of antibodies that can specifically detect the protein of interest. Alternatively, there have been developed reporter *Drosophila* lines that express the protein of interest fused to a reporter protein that can be detected enzymatically or by fluorescence (Timmons *et al*. 1997; Chatterjee and Bohmann 2012). We have therefore used a transgenic *Drosophila* reporter line that carries two additional C-terminally tagged Sens-sfGFP reporters that can be detected either directly (live imaging) or by immunofluorescence (Sarov *et al*. 2016). We were therefore able to use two methods in order to examine Sens dynamics: direct detection of Sens using antibodies against the endogenous protein, and indirectly using anti-GFP antibodies. To validate that the reporter accurately reflects endogenous Sens protein pattern and level we performed a double staining experiment against endogenous Sens using an anti-Sens antibody (obtained from H. Bellen lab; Nolo et al., 2000), and an anti-sfGFP antibody, in both embryos and wing discs (Fig.2). Under these conditions, we expect that anti-Sens antibodies tag protein products deriving from 4 *Senseless* genes (2 wild type and 2 sfGFP-tagged), while anti-sfGFP antibodies detect products from only the 2 sfGFP-tagged genes.

**Figure 2.**
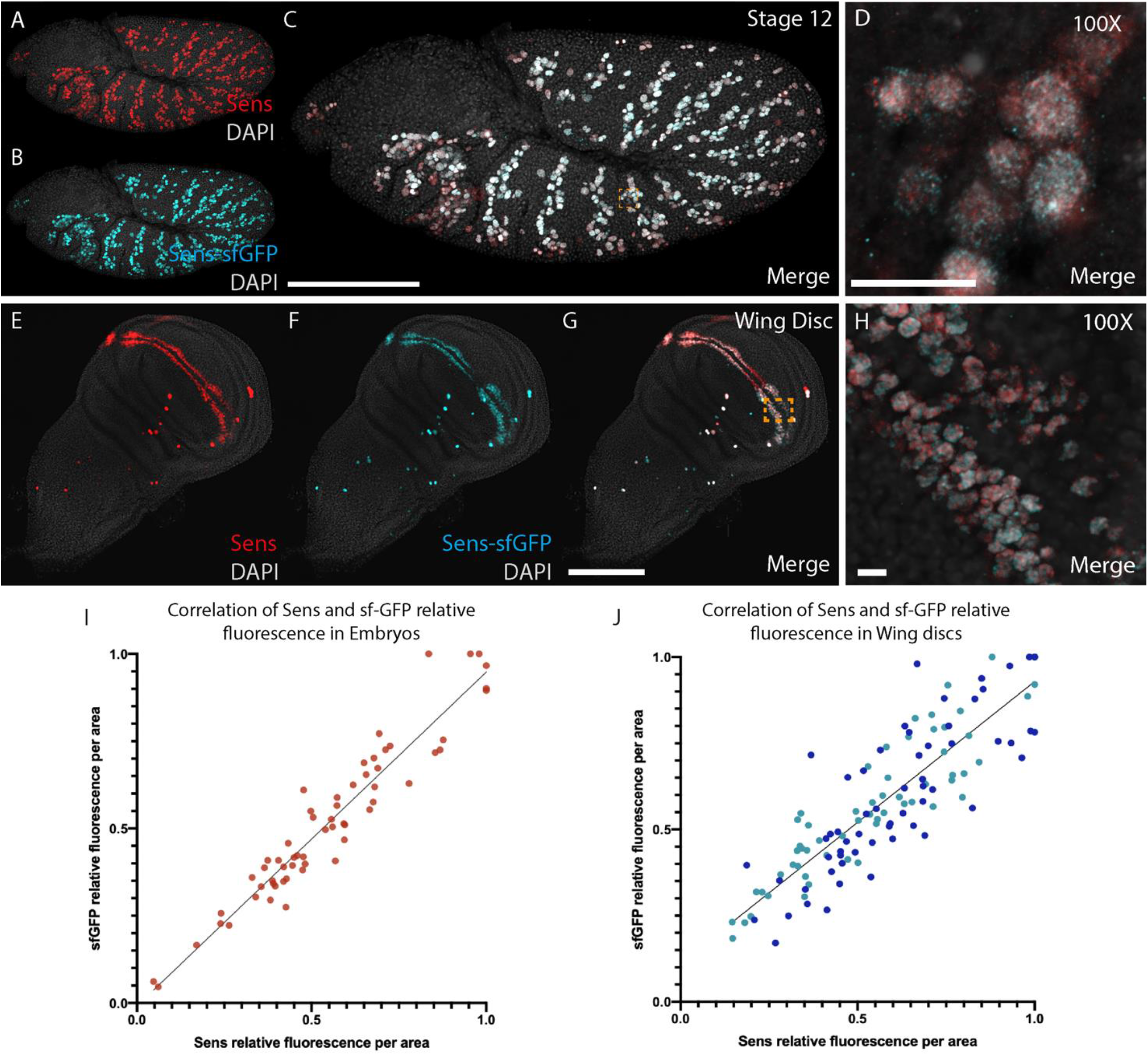
Colocalization of sfGFP reporter with endogenous Sens. Transgenic stage 12 embryo stained with antibodies against Sens (A, red) and sfGFP (B, cyan). The two antibodies colocalize (merged in D,E), showing that the sfGFP reporter is expressed in the same cells that are expressing endogenous Sens. Transgenic third instar larval wing disc stained with antibodies against Sens (E, red) and sfGFP (F, cyan). Again, the two antibodies colocalize in the same cells (merged in G,H). Correlation between relative fluorescence of Sens antibody and sfGFP antibody in embryos (I) and wing disc (J). For each of 3 embryos, fluorescence measurements were performed in 10 cells, while for each of 3 wing disc, 20 cells were measured. For each replicate, antibody fluorescence was normalized using the maximum value measured in that replicate. Scalebars: (C) 100 μm, (D) 10 μm, (G) 100 μm, (H) 10 μm.

The co-localization of the WT and sfGFP-tagged signals in embryogenesis and wing disc development are shown in Fig. 2. The data clearly show that during both embryogenesis (Fig. 2 A-D) and wing disc development (Fig 2 E-H), the two signals co-localize in the same cells. We are therefore confident that the reporter accurately recapitulates the endogenous pattern of Sens expression during embryonic and wing imaginal disc development. We further measured the relative fluorescence from each of the two antibodies in single cells to ensure that the reporter gene reflects Sens abundance. As expected, the sfGFP reporter is expressed proportionally to *Senseless* in both embryos (Fig. 2 D) and wing discs (Fig. 2 I). This indicates that the fluorescence signal from antibodies against sfGFP provides reliable information on endogenous *Senseless* localization and relative expression levels.

### *mir-9a expression pattern* and *Senseless* protein levels are inversely correlated during embryogenesis

In order to study the reciprocal dynamics of *mir-9a* and *Senseless* during embryogenesis and wing disc development, we simultaneously tracked active sites of *mir-9a* transcription (transcription sites – TS) using smFISH and Sens abundance via IF. We investigated these patterns at three different stages of embryonic development: stage 10, 11 and 12 (Fig. 3 A-C) after early *Senseless* expression and the initial stages of SOP specification. Interestingly we found that *mir-9a* transcription levels were inversely corelated with *Senseless* protein levels, and that *mir-9a* transcription is repressed rapidly after the initiation of Sens expression. Intriguingly, we noticed that a very small number of *Senseless*-expressing cells also displayed active *mir-9a* TSs (Fig. 3 A’’-C’’). Moreover, when we look at the fluorescence levels of sfGFP and the size of *mir-9a* TSs in the subset of cells that express both, it is evident that both miRNA active transcription and Sens level are lower in comparison to the rest of the cells. We believe that *Senseless* has just started to be translated in these cells and *mir-9a* transcription is stopping. Our interpretation is that the accumulation of Sens in the nucleus is associated with repression of *mir-9a* transcription, either by direct repression, or through an intermediary negative regulator.

**Figure 3.**
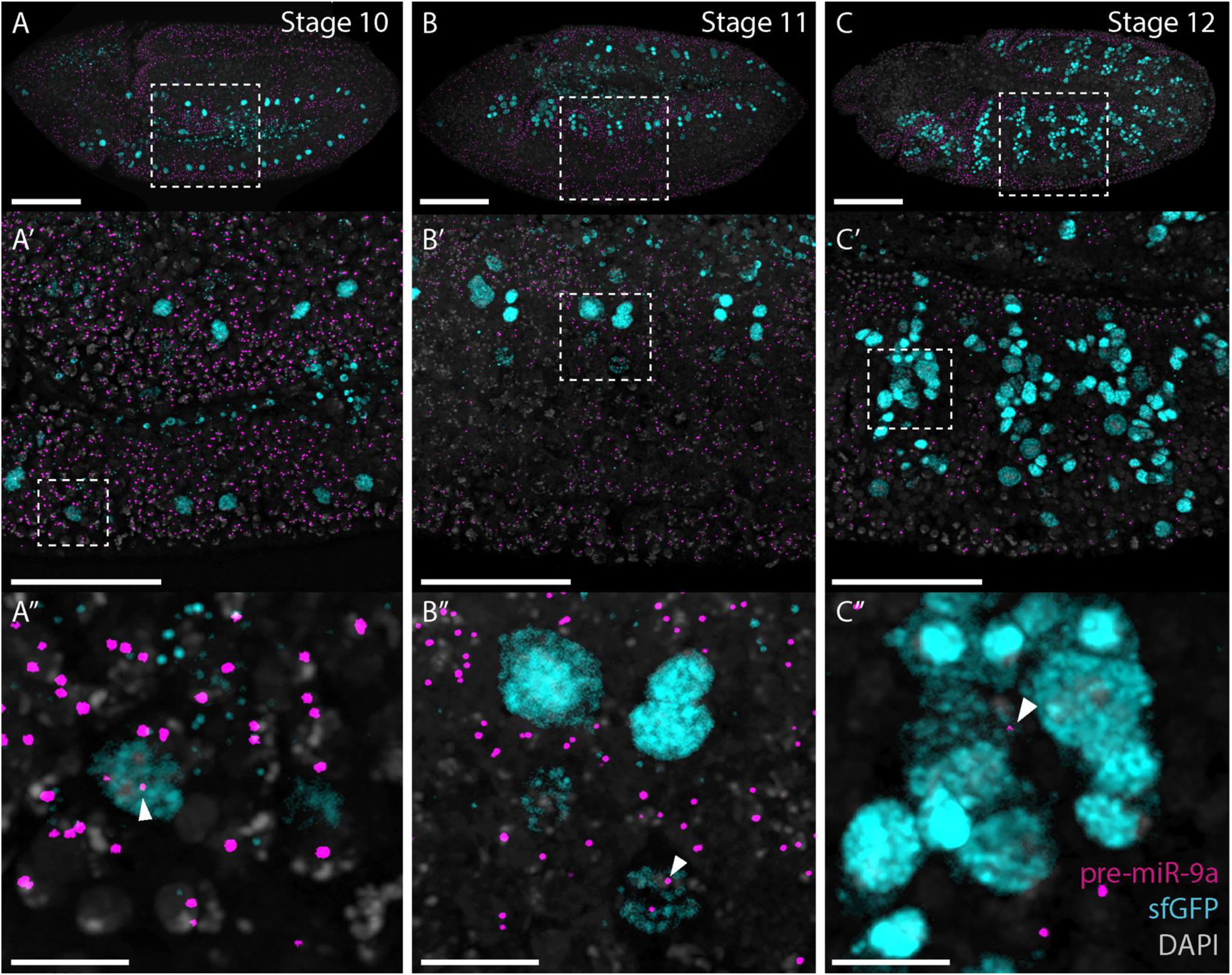
*mir-9a* is generally not actively transcribed in *Senseless* expressing cells during embryogenesis. (A-C) Transgenic embryos stained with probes against *pri-mir-9a* (magenta) and antibodies against sfGFP (cyan). (A,A’,A’’) Stage 10 embryo. Sens is expressed in isolated cells. Some Sens-expressing cells have active sites of transcription for *mir-9a*, but these appear to be less intense. (B,B’,B’’) Stage 11 embryo. Sens is expressed in more cells, a few of which still transcribe *mir-9a*. (C,C’,C’’) Stage 12 embryo. Sens expression reaches its peak during embryogenesis and *mir-9a* is generally not transcribed in Sens-expressing cells. Scalebars: (A-C) 100 μm, (A’-C’) 50 μm, (A’’-C’’) 10 μm.

To further investigate the dynamic relationship between expression of *mir-9a* and Sens, we developed a multi-channel experiment to simultaneously study the expression pattern of *mir-9a* TSs and Sens-sfGFP with *Senseless* and sfGFP mRNAs (Fig. 4). Intriguingly, we observe that cells that are transcribing *Senseless*, but that have not yet accumulated observable quantities of Sens protein (total lack of sfGFP signal), transcribe *mir-9a* (Fig. 4 B). *Senseless* expressing cells occupy several embryonic cellular layers. From an orthogonal projection that allows clear visualization of embryonic cell layers, we can see that cells containing Sens protein are located inwards, whereas cells that are transcribing both *mir-9a* and *Senseless* mRNA, but not yet translating Sens protein, are usually localized on the embryonic surface (Fig.4 C). We believe that the rapid dynamic changes in *mir-9a* expression are correlated with SOP differentiation, during which SOPs progressively migrate inwards as Sens protein accumulates and *mir-9a* is turned off.

**Figure 4.**
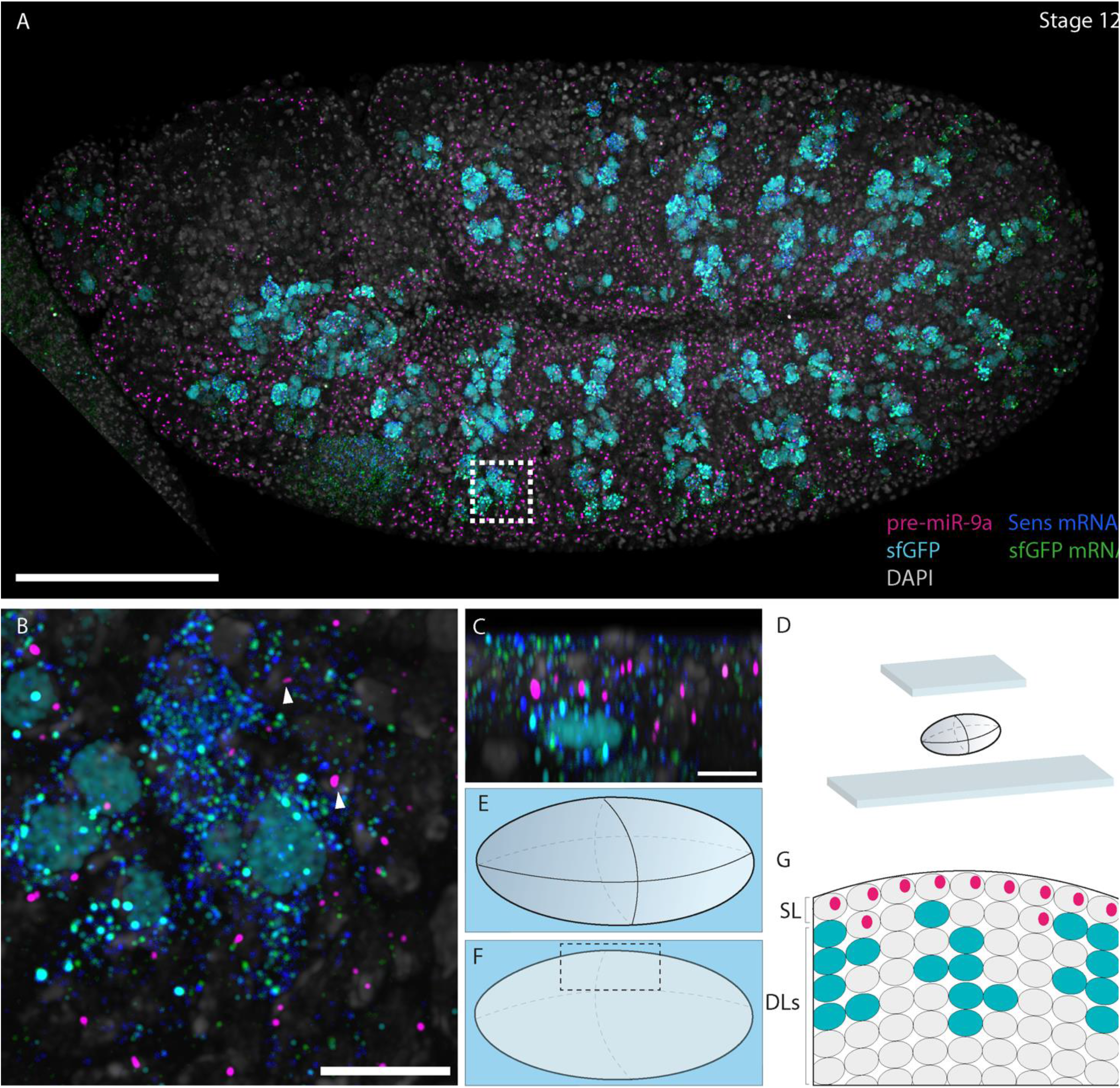
*miR-9a* is actively transcribed after *Senseless* transcription starts but before Sens protein is detectable. (A) Stage 12 transgenic embryo stained with probes against pri-*mir-9a*, sfGFP mRNA and *Senseless* mRNA and antibody against sfGFP. (B) zoom from different focal plane showing that some cells transcribing *Senseless* are still transcribing *mir-9a*. (C) Orthogonal view from (B) showing *mir-9a* is mostly expressed in cells at the embryo surface, some of which show detectable *Senseless* and GFP mRNA. Cells that already have Sens protein have migrated inwards. (D) Schematic representation of embryo mounting after smFISH and/or IF. (E) Lateral view of an embryo. (F) Lateral section of embryo in (E). (G) Schematic representation of what is reported in panel (C). SL = superficial cellular layer, DLs = deeper cellular layers. Scalebars: (A) 100 μm, (B, C) 10 μm.

### *mir-9a* is actively transcribed in cells containing Sens during early SO specification in the third instar imaginal wing disc

Since *Senseless* regulates SOP differentiation during PNS development in the *Drosophila* larvae (Singhania and Grueber 2014), we applied the approach outlined above to study *mir-9a* expression pattern and Sens abundance in third instar imaginal wing discs at the single cell level. The adult *Drosophila* wing possesses a spatially organized series of bristles (a class of SO) located at the wing margin (Lu *et al*. 2011). Flies in which Sens is ectopically expressed in the wing disc exhibit an increased bristle number in that wing region (Jafar-Nejad *et al*. 2003). During larval development, *Senseless* starts to be expressed at around 15h after third instar ecdysis in single SOPs in the wing notum. At around 30h *Senseless* is expressed in an increased number of isolated SOPs in the wing and notum area plus in 2 distinct stripes of cells in the wing disc pouch (Mirth *et al*. 2009). Cells belonging to these 2 rows of *Senseless*-expressing cells will give rise to the adult wing margin bristles (Jafar-Nejad *et al*. 2006).

By nascent transcript smFISH, we observed that *mir-9a* is expressed in nearly all cells in the wing disc. When we correlated *mir-9a* expression with that of *Senseless* in third instar discs we identified a small population of *Senseless* expressing cells with no *mir-9a* expression. These cells always had high levels of Sens protein, similar to our observations in the embryo. We also observed that many cells that have low or intermediate Sens protein levels are actively transcribing *mir-9a* (Fig. 5 A-B). It is interesting to note that *mir-9a* TSs size in *Senseless*-expressing cells do not differ from the size of TSs belonging to cells that are not expressing Sens protein. This indicates that these cells may not shut down *mir-9a* transcription, which might be kept active or modulated via transcriptional bursting. Nonetheless, at this stage only a minority of cells that contain Sens protein are not transcribing *mir-9a*. We therefore measured the intensity coming from sfGFP antibody (a proxy for Sens protein levels) at the single cell level to see if there was a difference in *Senseless* levels between *mir-9a*-expressing and non-expressing cells. The data clearly show that Sens is more abundant in cells that are not transcribing *mir-9a* (Fig. 5 C). However, our finding of concurrent expression of Sens and *mir-9a* contradicts aspects of the previously established model of triple row bristle specification (Li *et al*. 2006), which suggested a binary co-expression pattern.

**Figure 5.**
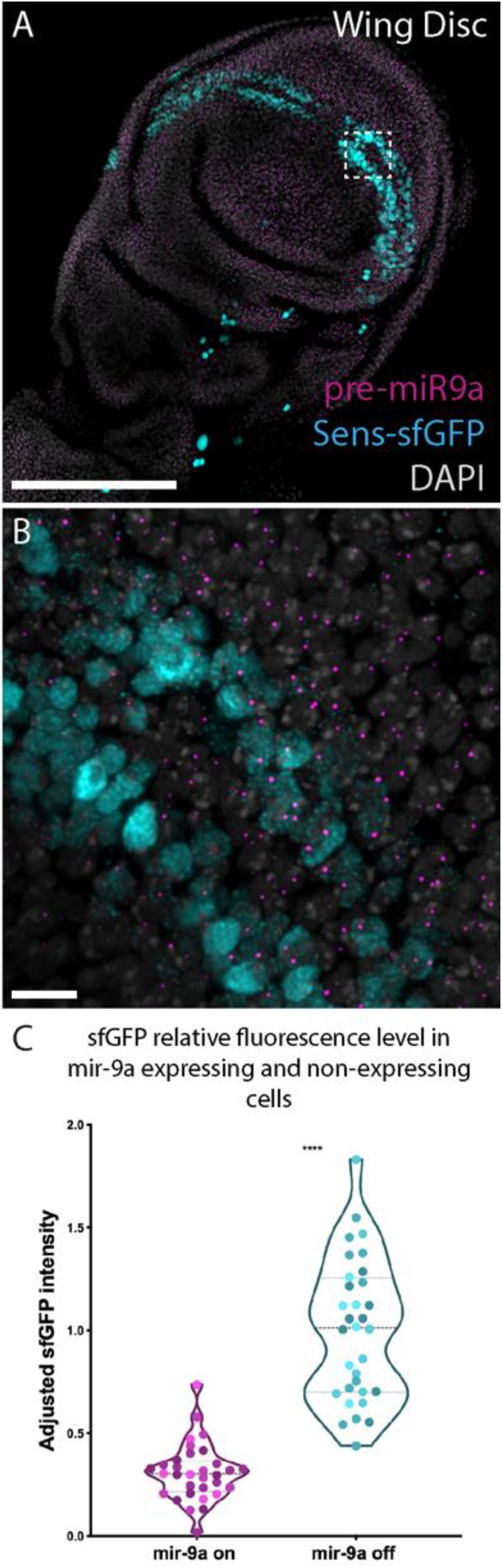
In the wing disc, Sens-expressing cells are generally actively transcribing *mir-9a*. (A-B) Third instar larval transgenic wing disc stained with probes against *pri-mir-9a* and antibody against sfGFP. Many cells that are actively transcribing *mir-9a* are also expressing Sens. (C) Adjusted sfGFP intensity coming from Sens expressing cells that are and are not actively transcribing *mir-9a*. Different colours represent data coming from different discs (n=4). Data from each replicate was normalized using the maximum adjusted fluorescence value from the group *mir-9a* off from that replicate. Scalebars: (A) 100μm, (B) 10 μm.

## Discussion

The stable and reproducible development of the *Drosophila* PNS is an extraordinary model of how the stereotyped stability of cellular differentiation is achieved (Jan and Jan 1994; Barad *et al*. 2011). In this study we focused on the role of *Drosophila miR-9a* in regulating the function of *Sens*, a crucial transcription factor that orchestrates SOP differentiation and PNS development in embryos and larvae. Dysregulated *miR-9a* expression results in disrupted *Sens* function leading to altered SOP differentiation and loss of stable stereotyped neural development (Li *et al*. 2006; Cassidy *et al*. 2013). It has been hypothesized that Notch signalling plays a key role in regulating *mir-9a* transcription in epithelial cells adjacent to potential SOPs thus preventing accumulation of Sens and unintended differentiation of additional SOPs (Li *et al*. 2006). Despite extensive study of *Sens* expression (Nolo *et al*. 2000; Mirth *et al*. 2009), there is little information regarding the developmental profile of *Sens* and *mir-9a* co-expression.

During embryonic development, we show that *mir-9a* is initially expressed throughout the neurogenic ectoderm, and a mutually exclusive expression pattern with *Sens* is established. Our single cell analysis shows that cells just initiating *Sens* expression, who therefore have not accumulated measurable Sens protein, actively transcribe *mir-9a*. However once Sens protein levels increase, *mir-9a* transcription is lost. The data suggest that *mir-9a* expression is turned off when the level of Sens protein reaches a specific threshold. Without *miR-9a* repression, Sens protein then accumulates rapidly, leading to SOP differentiation (Fig. 6 A-B). We suggest that this negative feedback loop involving Sens and *miR-9a* may be key in the regulative establishment of the SOP pattern. It is currently unclear if *Senseless* directly switches off *mir-9a* transcription or if it is an indirect effect of a multi-level genetic cascade.

**Figure 6.**
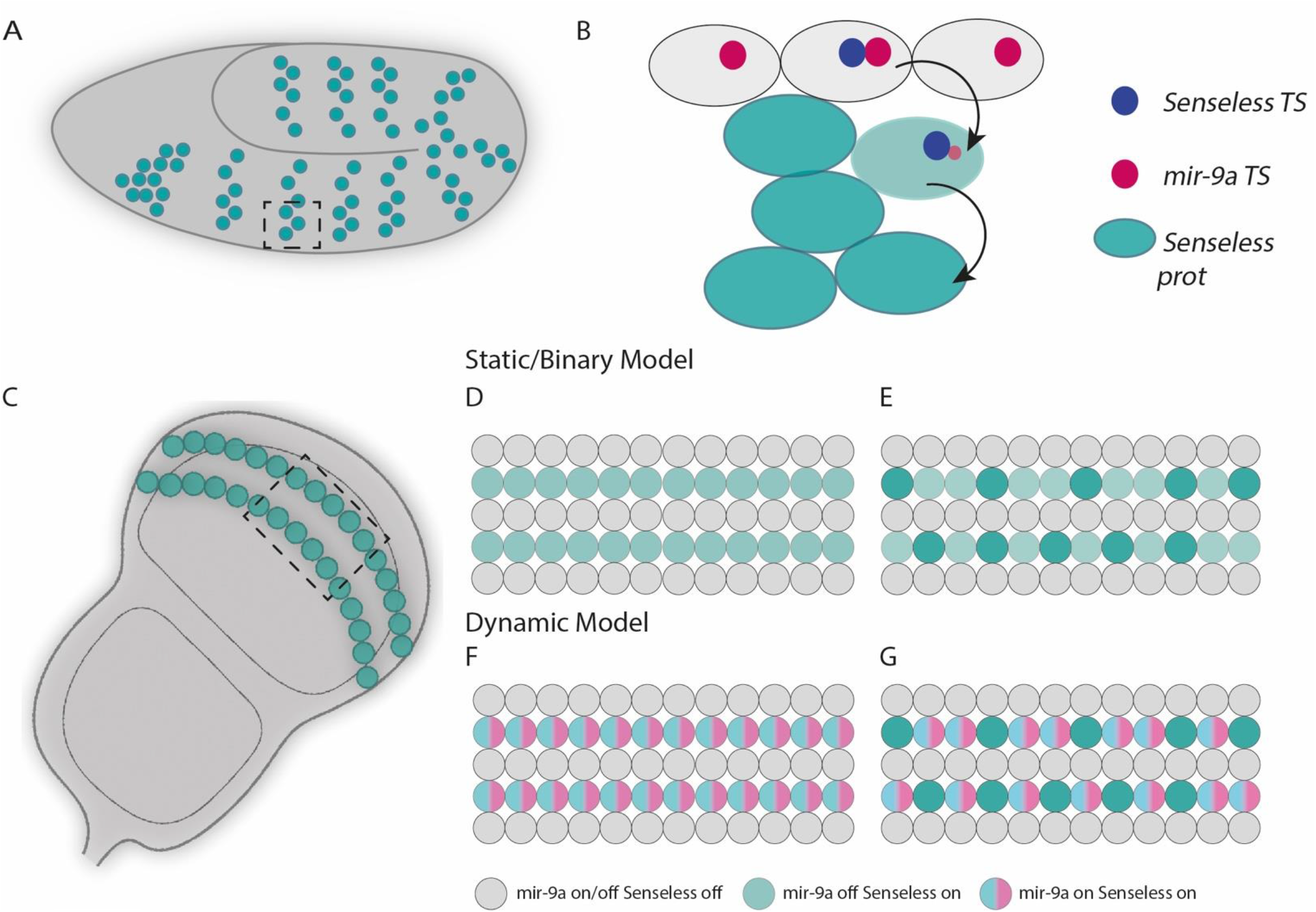
Model of *mir-9a* and *Senseless* interaction during embryogenesis and wing disc development. In the embryo (A) *Senseless* is expressed in clusters of cells. The orthogonal view of one of these clusters (B) shows that *mir-9a* is transcribed in cells at the top, some of which are turning *Senseless* transcription on. As *Senseless* mRNA gets translated, these cells stop transcribing *mir-9a* and move inwards. (C) In the *Drosophila* wing disc, *Senseless* is expressed in 2 rows of cells (plus additional isolated cells not shown). (D-E) In a static model, *mir-9a* and *Senseless* are not co-expressed. (F-G) Dynamic model in which all the cells that contain Sens are transcribing mir-9a, which is then turned off in a specific class of SOPs.

Li and co-workers presented evidence that in the third instar wing imaginal disc cells that express *Senseless* do not express *mir-9a*, which is otherwise widely expressed throughout the disc (Li *et al*. 2006). We find that *mir-9a* and *Senseless* are often though not always co-expressed in the wing disc. More specifically we find that among the *Senseless* expressing cells, those that are also transcribing *mir-9a* present a lower level of nuclear Sens, similar to that seen fleetingly prior to the establishment of the terminal and mutually exclusive pattern of SOP specification in the early embryo. The main difference here is that this subset of cells in the wing disc do not seem to be stopping *mir-9a* expression as it was happening in the embryo. This suggests that *mir-9a* and *Senseless* have an intricated reciprocal dynamic expression during embryogenesis and larval development.

Our findings complement the model (Fig. 6 D-E) presented by Li (Li *et al*. 2006) and show that *mir-9a* and *Senseless* exhibit dynamic co-expression rather than a binary one. Our findings are also in concordance with the suggestion by Jafar-Nejad (2006) that the genes that orchestrate PNS development in embryos and larvae might be different, even though the process follows a similar pattern. For instance, during embryogenesis *achete* and *scute* are necessary for *Sens* activation, while during larval development this function is accomplished by *Wingless* (Jafar-Nejad *et al*. 2006). *mir-9a* function might therefore differ between embryonic and larval SOP development via the presence or absence of other *mir-9a* targets. We propose that *mir-9a* repression during embryogenesis allows *Sens* to reach a specific threshold in order to establish the correct number and pattern of embryonic SOPs.

During larval development, *Sens* might be expressed at different levels depending on the subclass of SO and this in part involves regulatory feedback by *mir-9a*. The fly wing margin possesses 2 different kinds of SO, the mechanoreceptors and chemoreceptors, and these have a very precise pattern (Hartenstein 1993; Raad *et al*. 2016). *mir-9a* expression may be involved in a regulatory loop with *Sens* to set different *Sens* levels and thereby control the kind of SO that will develop from each particular SOP. Our data further suggest that *Sens*-expressing cells that are not transcribing *mir-9a* will adopt chemoreceptor SOP fate: chemoreceptors are lower in number than mechanoreceptors and their localization and alternation resembles the pattern of cells with high Sens expression level. Therefore, we believe that *mir-9a* serves to keep Sens expression low in mechanoreceptor precursor cells to ensure they adopt the correct SOP cell fate. Temporally, our model suggests that *mir-9a* is initially expressed in all *Sens*-expressing cells, delaying the adoption of a specific SOP fate (Fig. 6 F-G). As SOPs adopt their specific cell fate, *mir-9a* is switched off as a consequence of the transition from a multipotent precursor to a determined cell.

Our work suggests that *mir-9a* has a dynamic role in the specification of SOP differentiation, tuning *Sens* expression to ensure that the correct number of cells adopt the appropriate SOP fate. The molecular mechanism that dictates *mir-9a* transcription during PNS development remains unknown; its elucidation is important for complete understanding of this dynamic process. A fundamental question that needs to be answered is the mechanism by which the mutual regulation of *mir-9a* and *Sens* act to establish the observed co-expression dynamic, switching from mutually exclusive to co-expressed depending on the fly developmental stage. This study demonstrates the importance of examining miRNA regulation and miRNA-target gene expression dynamics at a single cell level in the developing organism. We suggest that these dynamic co-regulatory processes are a general feature of microRNA function during development.

## Acknowledgments

We thank the staff from the University of Manchester’ Bioimaging Facility, in particular Dr. Peter March, for help with confocal microscopy. We also thank Hugo J. Bellen from the Howard Hughes Medical Institute for providing the anti-Sens antibody and Dr. Fabian Morales-Polanco for discussions and comments on the manuscript. This work was funded by a Wellcome Trust funded 4-years PhD studentship [203808/Z/16/Z] to LG.

## Author contributions

LG, MR and SGJ conceived the project. Experiments were designed by LG and MR and performed by LG. The manuscript was written by all the authors.

## Supplemental Table 1

**Table 1.**
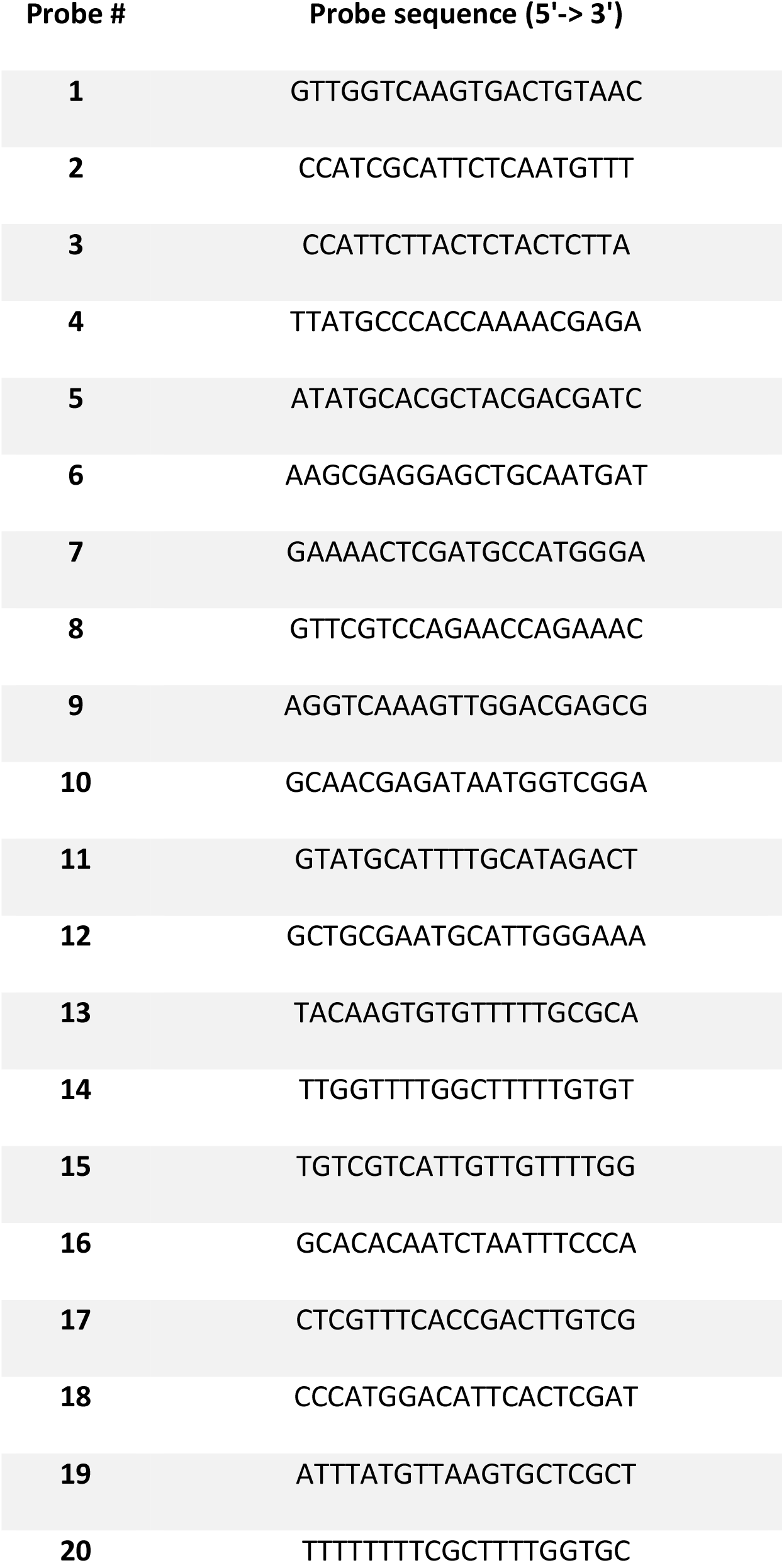
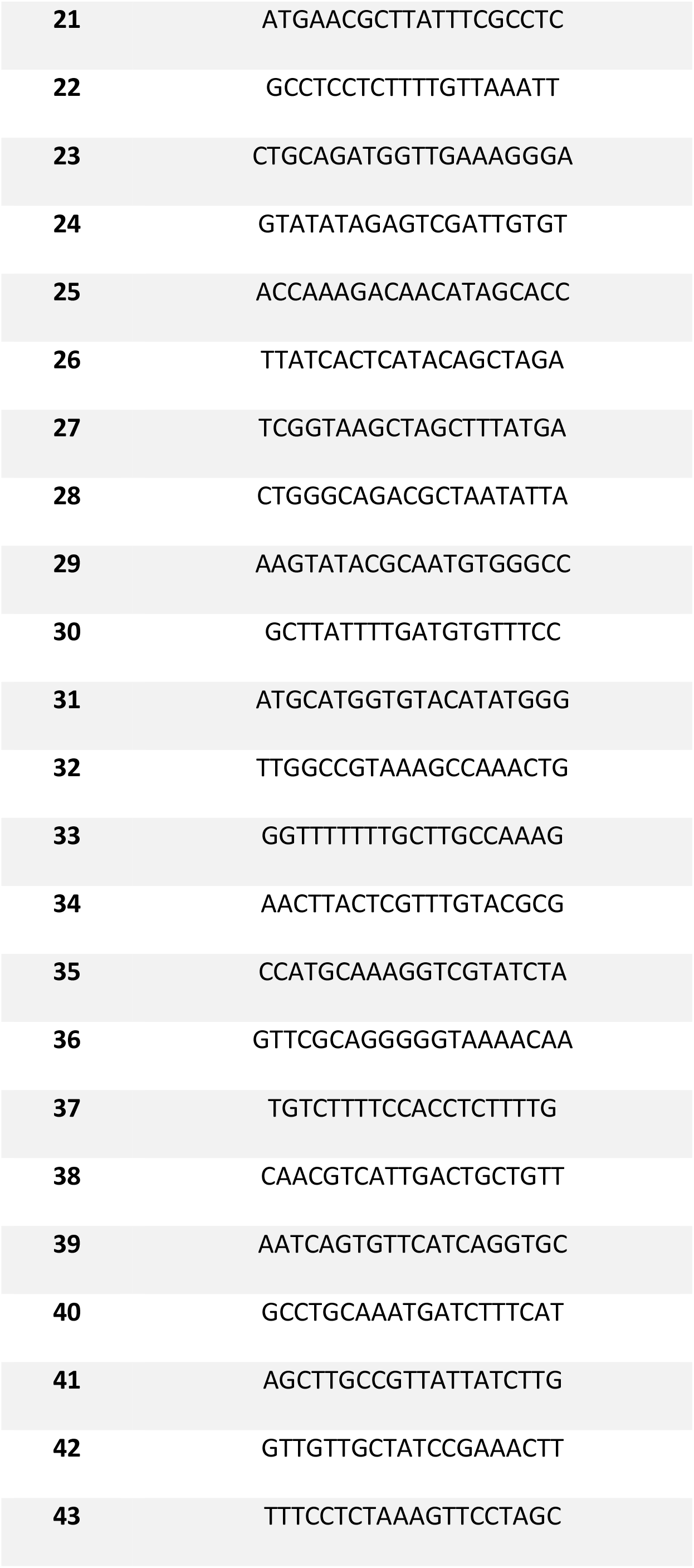
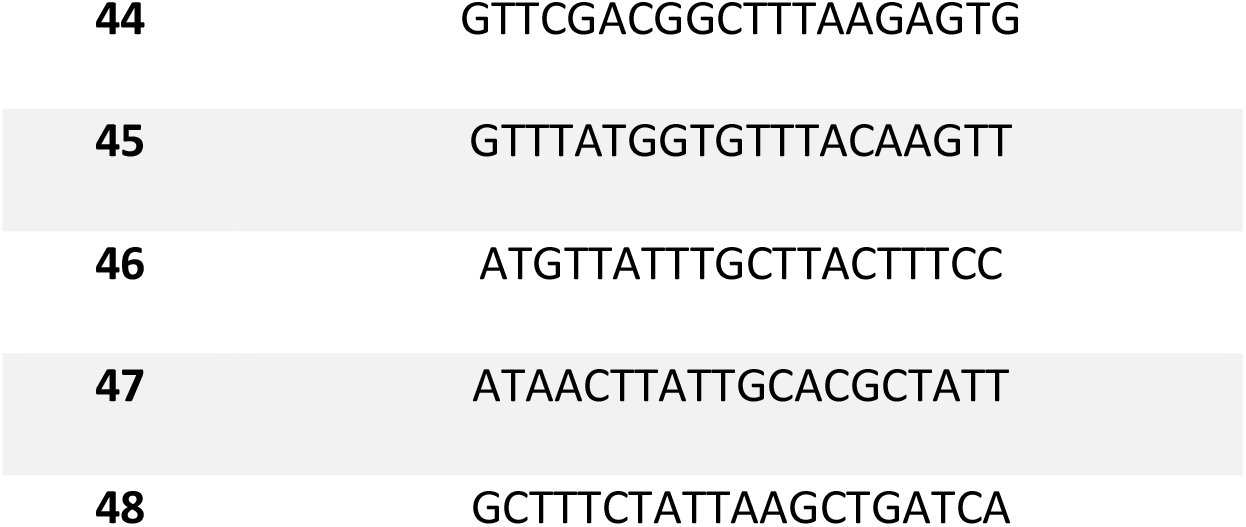
Probes against *pri-mir-9a*.

**Table 2.**
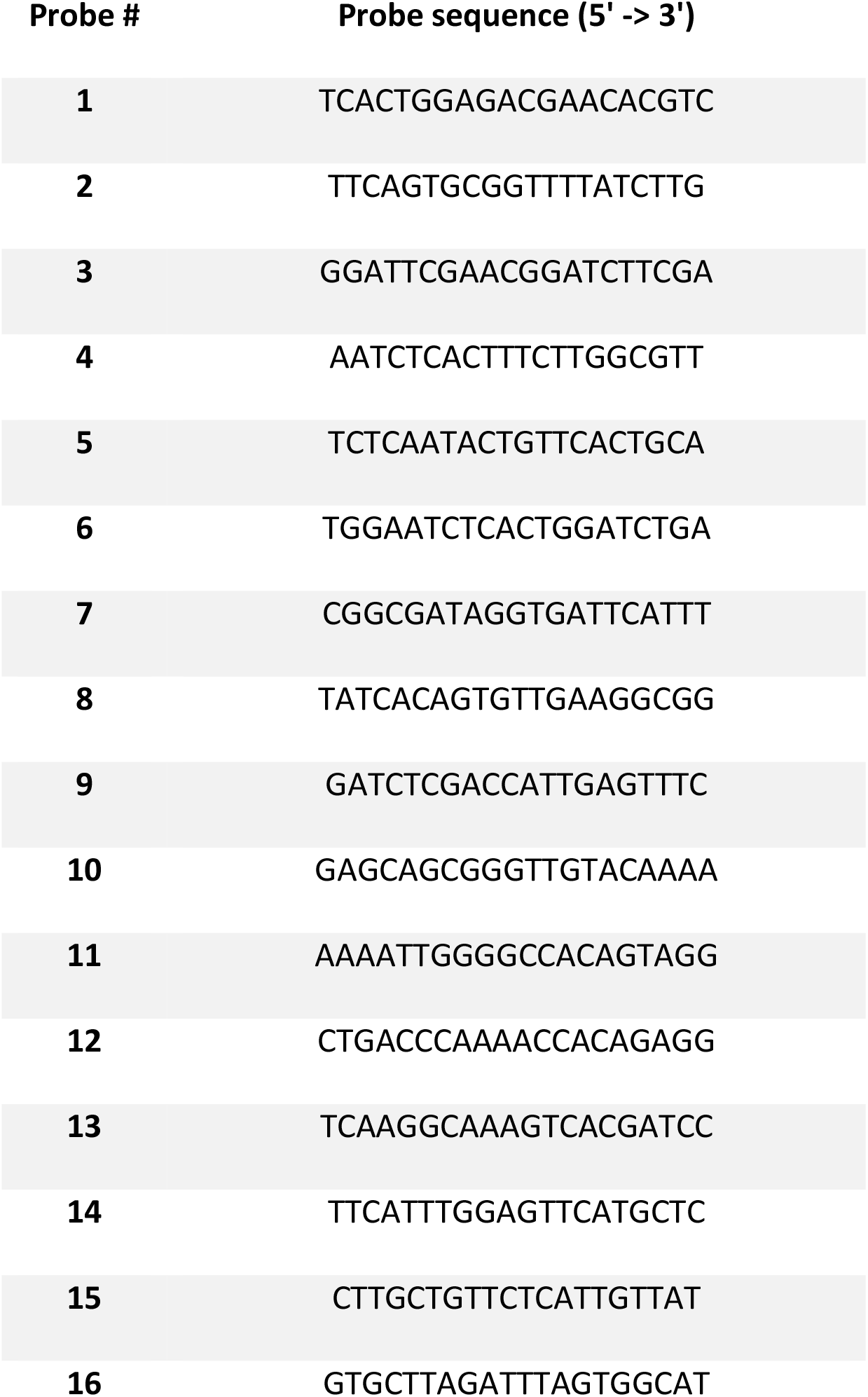
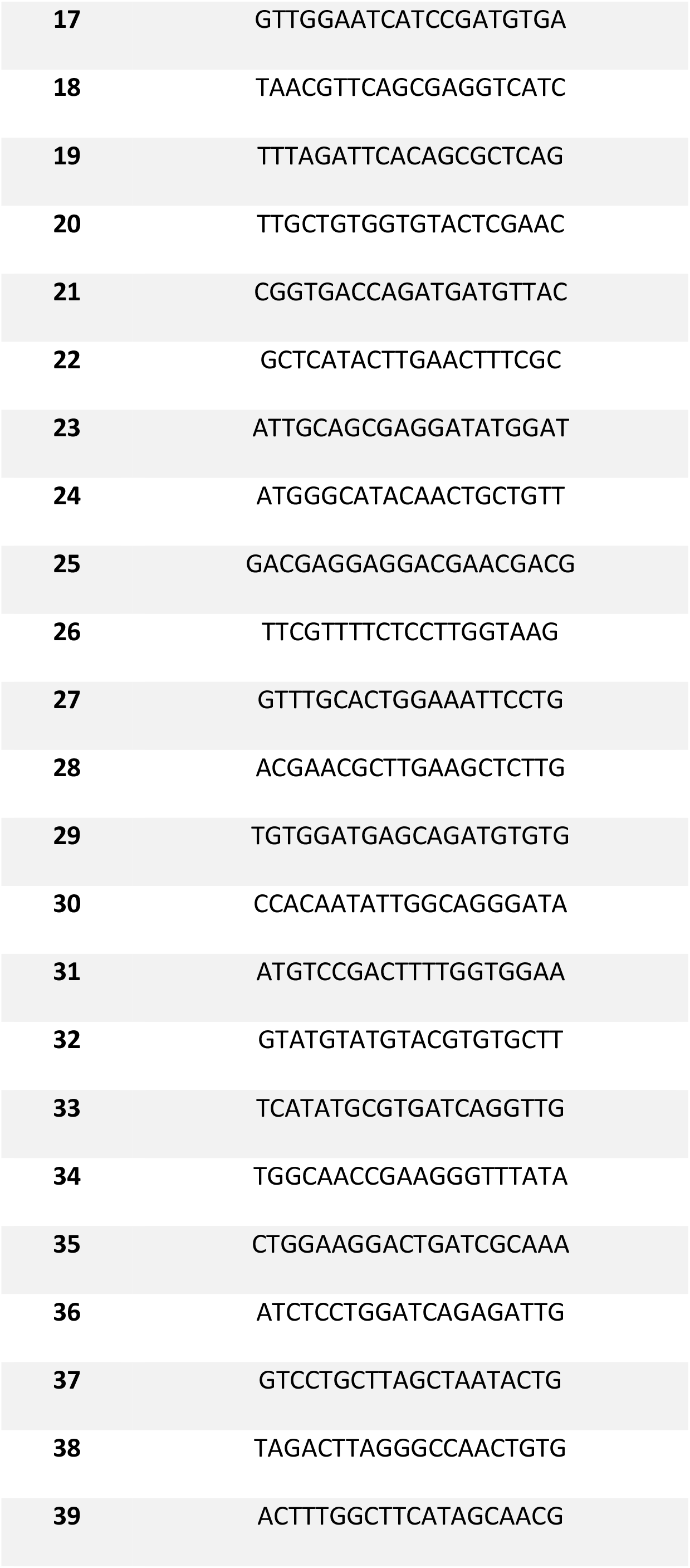
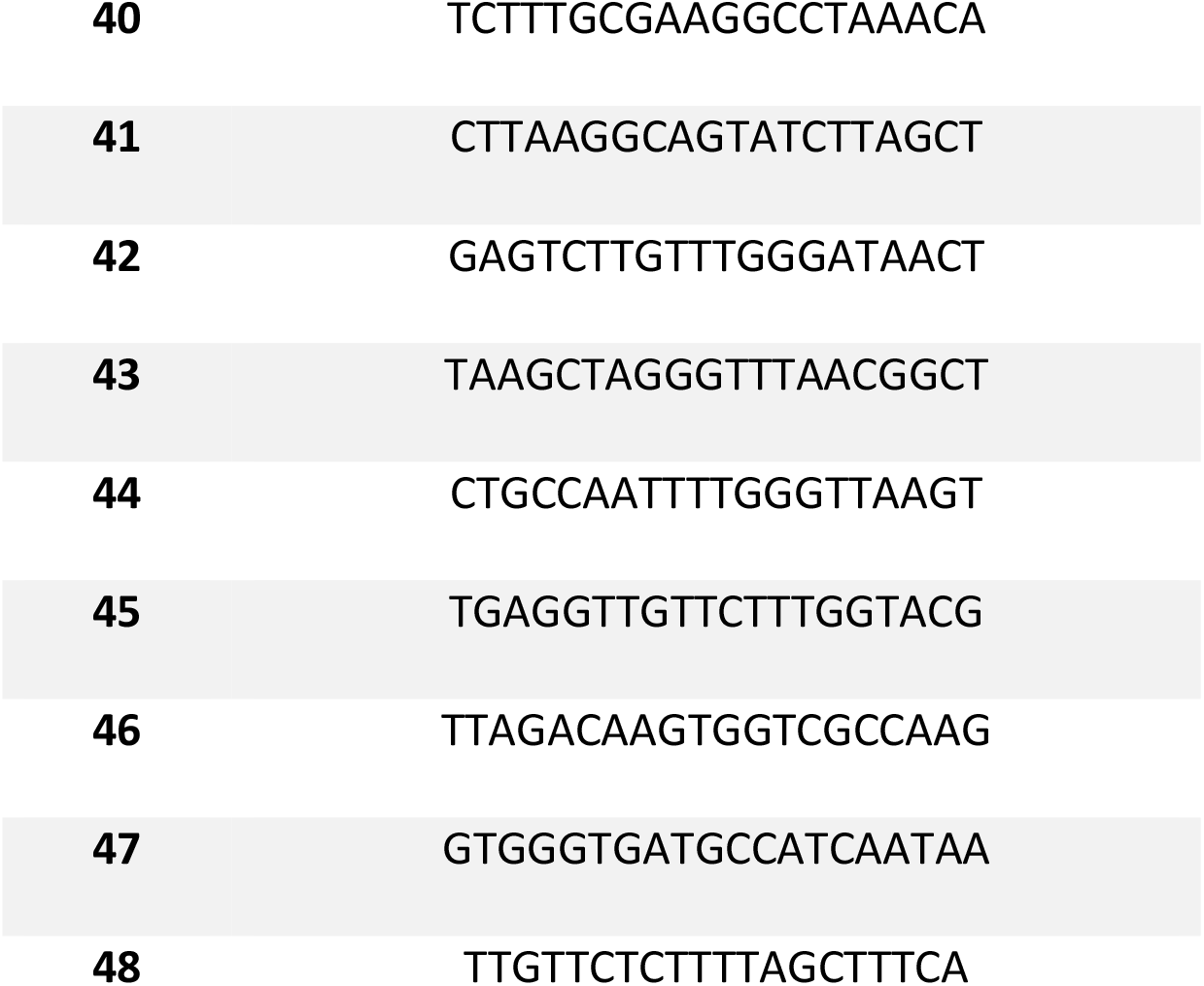
Probes against *Senseless*

**Table 3.**
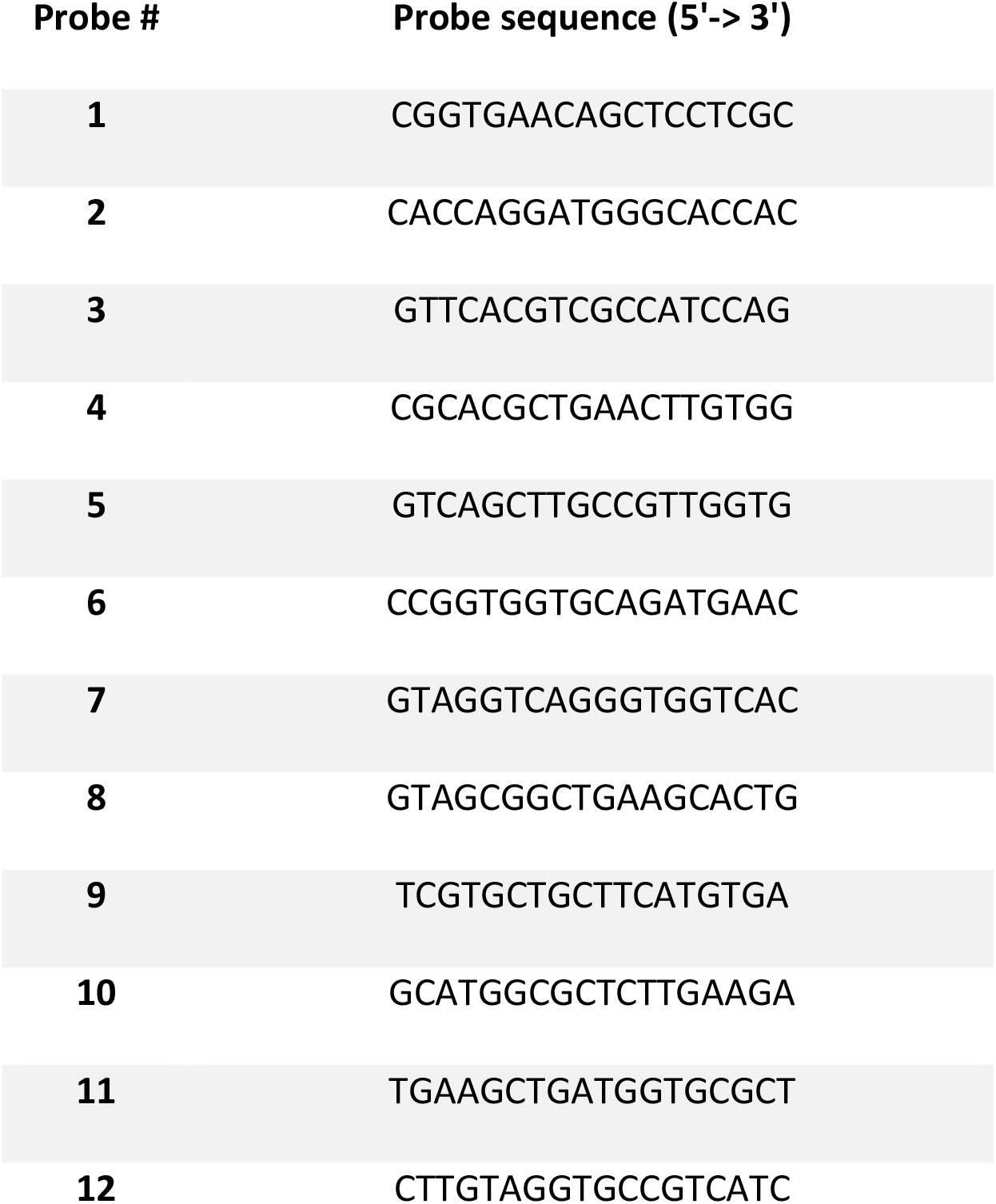
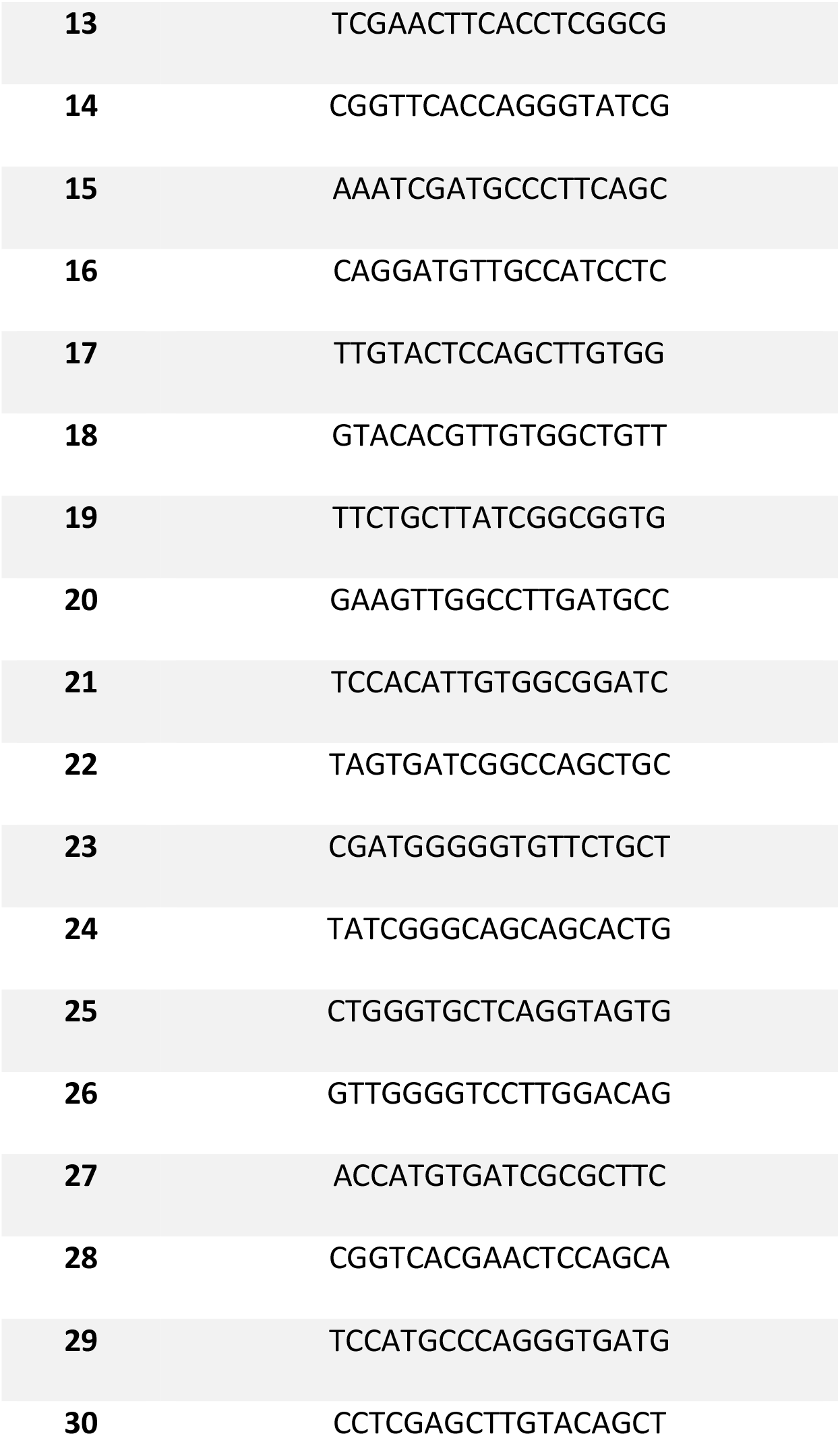
Probes against sfGFP.

---

**Supplementary Figure S1.**
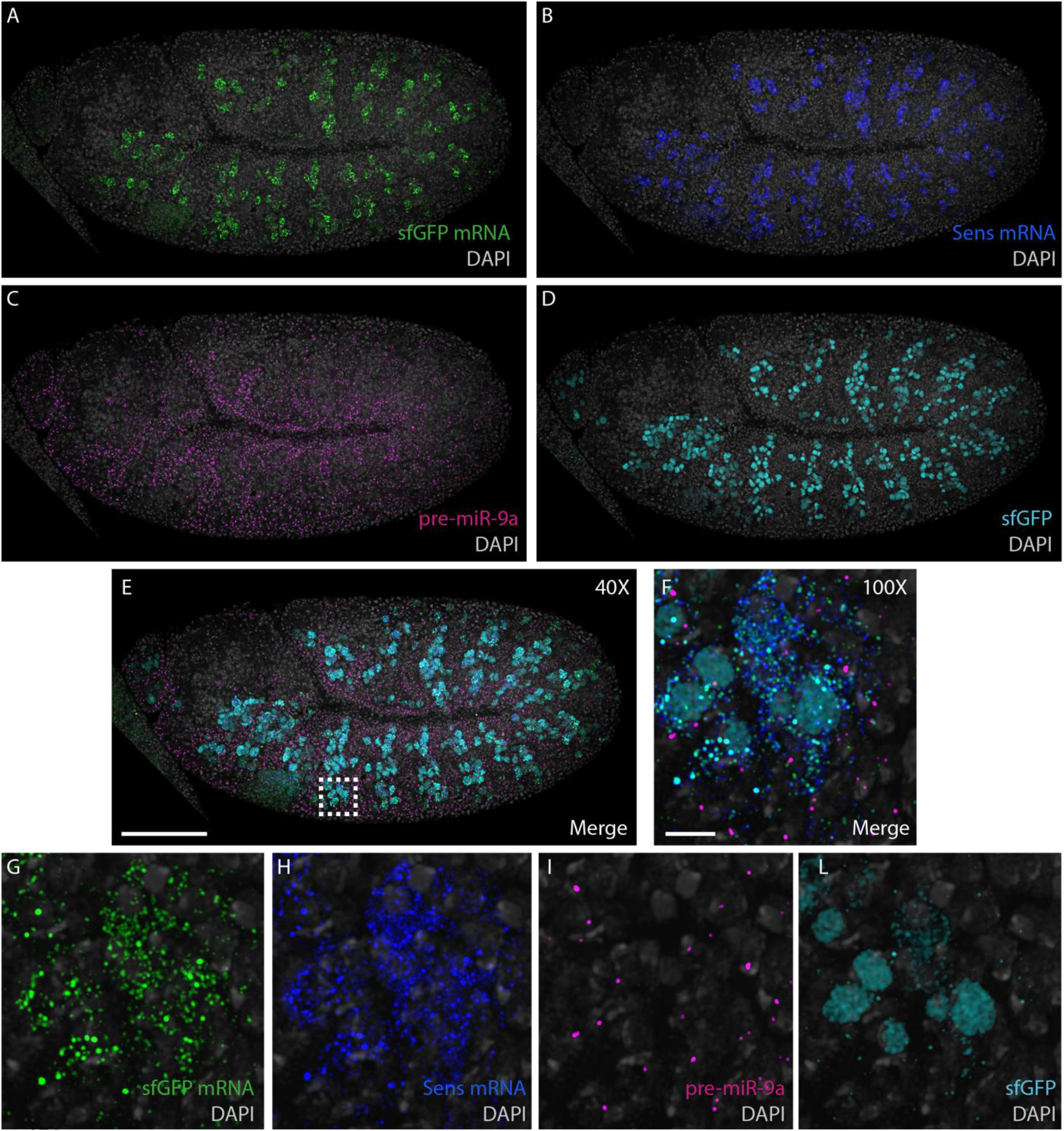
Single Channels from Fig.4 (A) sfGFP probes 40X. (B) *Senseless* probes 40X. (C) *pre-miR-9a* probes 40X. (D) sfGFP antibody 40X. (E) Marge 40X. (F) Merge 100X. (G) sfGFP probes 100X. (H) *Senseless* probes 100X. (I) *pre-miR-9a* probes 100X. (L) sfGFP antibody 100X.

